# Recruitment of the outer-membrane lipoprotein DolP to the division site via cardiolipin-mediated diffusion-state switching

**DOI:** 10.1101/2025.10.29.685289

**Authors:** Jinchan Xue, Di Yan, Jiajia Wang, Yuzhi Hong, Xinxing Yang

## Abstract

In diderm bacteria, the outer membrane (OM) must invaginate in step with septal peptidoglycan (PG) remodeling during cytokinesis. One OM lipoprotein, DolP, has been shown to localize at the cell division site and to facilitate the daughter cell separation. Yet how DolP is recruited to the division site remains unclear at the molecular level. Here, we show that DolP arrives concomitantly with the late divisome protein FtsN. Utilizing single-particle tracking Photoactivated Localization Microscopy (spt-PALM), we investigated the dynamics of individual DolP molecules in living *Escherichia coli* cells. Single-molecule analysis revealed two diffusion states: a diffusive state across the cell envelope and a previously underappreciated immobile state enriched at the septal and polar regions. Importantly, DolP does not comigrate with the processive FtsW-FtsI-FtsN complex. Mutations of the cardiolipin-binding surface caused loss of mid-cell enrichment and reduced the immobile fraction. Cardiolipin’s preference for high negative curvature enriches it at the division site, creating a trapping environment for DolP. Additionally, the active septal constriction but not the late divisome proteins recruit DolP. Together, these findings indicate that cardiolipin-mediated immobilization underlies DolP’s septal enrichment.

## Introduction

Cytokinesis demands the coordinated constriction of all three envelope layers in diderm bacterial cells—the inner membrane (IM), the peptidoglycan (PG) cell wall, and the outer membrane (OM). Recent works have made clear that the OM is not a passive passenger^1,2^: it contributes substantially to the envelope’s load-bearing capacity and shape stability^3^, and its invagination and fusion are essential for completing a successful cell division^4^. Perturbing OM stiffness or its connections to PG causes envelope deformation and compromises cell division under mechanical or osmotic stress^4,5^, underscoring the essential role of OM-related proteins in supporting PG synthesis-limited cell division^6^. The coupled invagination of the OM and IM with the septal PG has been illuminated through two complementary approaches. First, passive coupling via abundant OM–PG tethers (such as Lpp, OmpA, and PAL(peptidoglycan-associated lipoprotein)) can transmit inward motion as septal PG is synthesized, maintaining OM– PG spacing during constriction^7,8^. Second, active, energy-coupled constriction via the Tol–Pal system, in which the IM complex (TolQ– TolR–TolA) harnesses the proton motive force to engage TolB–Pal and promote OM invagination with IM at mid-cell^9^.

The invagination of septal PG relies on the PG hydrolases, mainly PG-related amidases. In *E. coli*, three autoinhibited amidases (AmiA, AmiB, AmiC) can cleave the peptide stems from the glycan strands and play an important role in cell separation. These enzymes are activated by two main regulatory pathways: FtsEX-EnvC and NlpD ^10^. From the IM side, the core divisome components FtsEX recruit EnvC that further activates AmiA and AmiB and feeds back to septal PG synthesis^11-14^. On the OM side, a lipoprotein NlpD activates AmiC to allow the inward PG degradation, thus coordinating the OM invagination with the septum constriction^4,5^.

Within this framework, the OM lipoprotein DolP (division and OM stress-associated lipid-binding protein; formerly YraP) has emerged as an OM-associated cell division protein. Genetic and physiological studies linked DolP to daughter-cell separation phenotypes that parallels to the EnvC-AmiAB pathway, although its precise function is still under debate^5,15^. Structural work showed that DolP is a dual BON (Bacterial OsmY and nodulation)-domain protein (Fig. 1A)^2,16^. Its C-terminal BON2 domain binds anionic phospholipids (notably cardiolipin, CL) through an extended surface centered on W127. The cardiolipin binding is a critical factor for DolP’s septal localization. Yet how DolP behaves biophysically in the OM and is recruited to the division site was not investigated.

**Figure 1.**
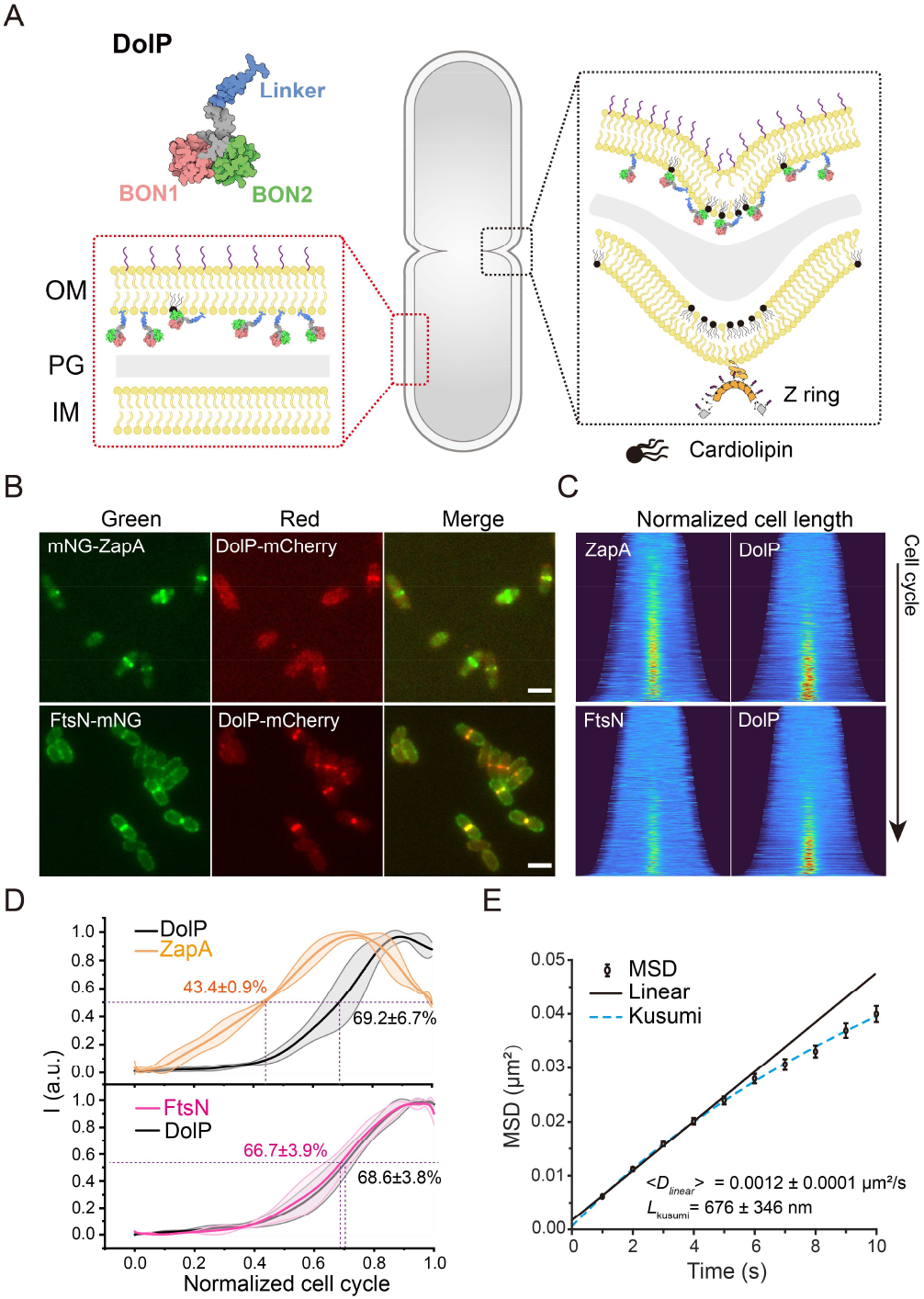
DolP is a late divisome protein but does not comigrate with septal PG synthases. **(A)** Schematic of DolP architecture (dual BON domains with a flexible linker connected to OM) and envelope context. Cardiolipin on the inner leaflet accumulates at regions of high negative curvature such as the constricting septum, where DolP is enriched. **(B)** Two-color imaging of cells expressing DolP–mCherry with mNeonGreen-ZapA (early divisome, top) or FtsN–mNeonGreen (late divisome, bottom). DolP colocalizes with FtsN but appears after ZapA. Scale bars, 2 µm. **(C)** Demographs of ZapA/FtsN (left of each panel pair) and DolP (right). More than 1000 cells from three independent experiments for each condition (Supplementary table 3). **(D)** Cell-cycle kinetics of mid-cell enrichment. Half-maximum times (mean ± s.d.) are labeled. **(E)** MSD curve of wildtype DolP versus time (black). Linear fit (solid line) and confined-diffusion (Kusumi) fit (dashed) yield ⟨*D*_linear_⟩ = 0.0012 ± 0.0001 µm^2.^s^−1^ (mean ± s.e.m.) with confinement length *L*_kusumi_ = 676 ± 346 nm (mean ± s.e.m.).

Here, by combining dual-color fluorescence imaging with Single-Particle Tracking Photoactivated Localization Microscopy (spt-PALM), we define DolP’s recruitment timing and mobility states in living *E. coli* cells. At the single-molecule level, DolP exists in two diffusion states on the inner leaflet of the OM—a previously unappreciated, nearly immobile state and a diffusive state. Importantly, we found that DolP’s recruitment depends on the immobile state enriched at the cardiolipin-rich, highly curved constriction zones but does not complex with the processive FtsWI septal PG synthase. These findings provide fundamental knowledge on the OM-lipoproteins’ mobility and explain the biophysical mechanism of how they sense membrane microenvironments to facilitate cytokinesis.

## Results

### DolP is recruited late to the divisome but does not comigrate with the core septal PG synthases

To establish when DolP is recruited at the division site, we expressed DolP–mCherry^17^ from an inducible plasmid. It has been shown that the cells lacking both *envC* and *dolP* genes exhibited a pronounced filamentation phenotype^5^. This DolP–mCherry fusion successfully complemented the filamentation phenotype of the Δ*envC*Δ*dolP* strain, indicating functionality (Fig. S1A,B; Supplementary Table 1,2). Across induction conditions, DolP-mCherry enriched at mid-cell in dividing cells (Fig. S1C), confirming it as a division-associated OM factor^5^.

We next compared the arrival time of DolP-mCherry using two-color imaging with mNeonGreen-ZapA^18^ (early divisome) and FtsN-mNeonGreen^19^ (late divisome)^20^. Demographs showed that DolP reached mid-cell after ZapA and approximately coincident with FtsN, at the onset of constriction (Fig. 1B,C; Supplementary Table 3). Quantitatively, mid-cell enrichment kinetics reached half-maximum at ∼69% of the cell cycle for DolP, compared with ∼43% for ZapA and ∼67% for FtsN (Fig. 1D), designating DolP as a late divisome factor. We then asked whether DolP physically associates with the canonical septal PG synthase complex, including FtsQLB, FtsWI, and FtsN, which move directionally around the septal ring during constriction^14,18,21^. Using PAmCherry^22^ fused DolP, we performed spt-PALM with 1-second exposure^23,24^ to favor detection of slow processive movement (Fig. S1A,B; Supplementary Table 6). However, no directional DolP motion was detected. Mean squared displacement (MSD) analysis indicated that DolP exhibited slow diffusion with slight confinement (average diffusion coefficient <*D*_linear_> =0.0012 ±0.0001 µm^2.^s^−1^, mean ±s.e.m.; confinement radius <*L*_kusumi_> =676 ±346 nm, mean ±s.e.m.; Fig. 1E). Short-exposure (50 ms) recordings likewise yielded slow diffusion without directional runs (Fig. S2A). Thus, DolP accumulates at mid-cell (the constriction site in *E. coli*) as constriction begins (with FtsN) but does not comigrate with FtsWI, implying exclusion from the core septal PG synthase complex.

### DolP exhibits two diffusion states and accumulates in the immobile state at division sites

Despite the absence of directional motion, single-molecule tracking revealed two mobility states of DolP. A subset of tracks remained confined/immobile over the observation window (Fig. 2A,B), whereas others diffused along the envelope (Fig. 2C,D). Cumulative radial distribution functions (CDFs) were well fit by a two-population model: an immobile state (*D*_1_ <1.0×10^−4^ µm^2.^s^−1^) and a diffusive state (*D*_2_ =0.0033 ±0.0002 µm^2.^s^−1^, mean ±s.e.m.) (Methods; Fig. 2E, Fig. S2B, Supplementary Table 5). The averaged cellular distribution of all DolP-PAmCherry localizations also displayed a mid-cell enrichment pattern, agreeing with the DolP-mCherry’s distribution (Methods; Fig. 2F).

**Figure 2.**
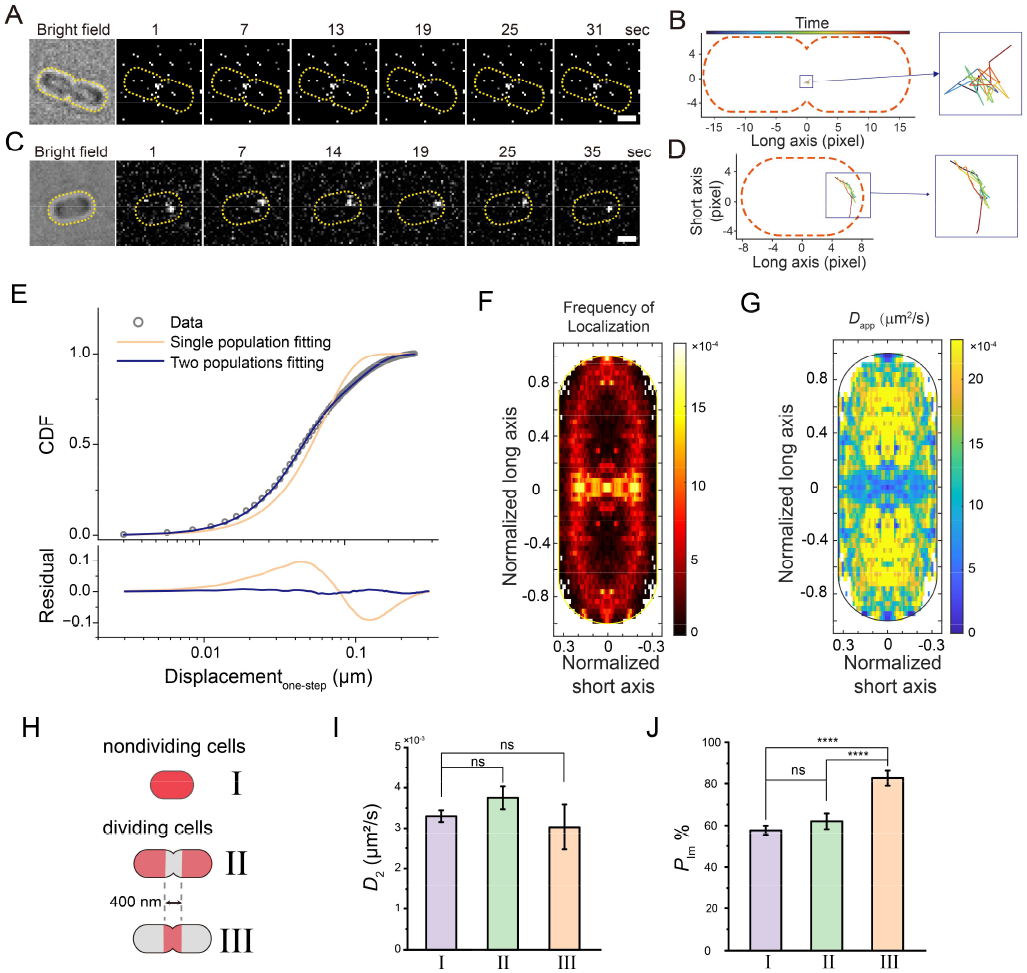
DolP exhibits two diffusion states and shifts toward an immobile state at the division site. **(A**,**C)** Montage images of two representative cells of single DolP molecules spt-PALM localizations for two representative cells. Scale bars, 1 µm. **(B**,**D)** Extracted trajectories from boxed regions in (A,C) illustrating confined/immobile and diffusive behaviors. **(E)** Cumulative radial distribution functions (CDFs) for one-step displacements. A two-population fit (blue) outperforms a single-population fit (tan) with residuals at the bottom. **(F)** Heatmap of DolP localization frequency in a normalized cell reveals mid-cell enrichment. **(G)** Heatmap of per-step apparent diffusion coefficient (*D*_app_) shows low-*D*_app_ enrichment at mid-cell (and poles). **(H)** Schematic of trajectory groups: trajectories in I, non-constricting cells; II, constricting cells, but off-septum; III, constricting cells within ±200 nm of mid-cell. **(I)** Diffusion coefficients of the diffusive state (*D*_2_) is similar across groups. **(J)** Immobile-state fraction (*P*_Im_) is significantly higher at constriction sites (class III ∼83%) than in classes II (∼62%) or I (∼58%). ****P < 0.0001; ns, not significant (P > 0.05); two-tailed Student’s t-test. More than 90 trajectories from three independent experiments for each subcellular group (Supplementary table 6).

To determine if the mobility of DolP is uniform across the cell, we computed the apparent single-step diffusion coefficient (*D*_app_) of DolP at each localization in a normalized cell (Methods; Fig. 2G). The heatmap showed pronounced enrichment of low *D*_app_ at mid-cell, mirroring the distribution of DolP itself. This distribution indicates that the more immobile DolP molecules are concentrated at the division site. Interestingly, DolP also displayed slow diffusion at the cell poles, where the OM is more negatively curved than the lateral regions of the cell.

We then categorized trajectories into three groups based on their localization and cell morphology: (I) in non-constricting cells, (II) in constricting cells but in off-septum regions, and (III) in constricting cells and within ±200 nm of the mid-cell (Fig. 2H). The group III trajectories represent DolP molecules in the division site whereas the group I and II DolP molecules do not associate with the divisome. Two-population CDF fits yielded similar *D* values across groups (*D*_1_ < 1.0×10^−4^ µm^2.^s^−1^, *D*_2_ ≈ 0.0030 µm^2.^s^−1^, Fig. 2I, Supplementary Table 6), but state occupancies differed: the immobile state accounted for ∼83 ±4% (mean ±s.e.m.) in group III versus ∼62% ±4% in group II and ∼58% ±2% in group I (Fig. 2J). MSD curves mirrored these occupancy changes, demonstrating reduced mobility of DolP at the division site (Fig. S2C). The enrichment of immobile state at mid-cell confirmed the results of the *D*_app_ heatmap and suggested DolP enrichment at mid-cell arises from a local shift in diffusion state toward an immobile/engaged population.

### Cardiolipin-binding site governs DolP’s diffusion and mid-cell retention

Given that cardiolipin accumulates at regions of negative membrane curvature^25^, and DolP’s BON2 domain binds anionic phospholipids, we reasoned that the cardiolipin-enriched microenvironment of the OM at septa may serve as the spatial cue and binding patterner that slows down DolP’s diffusion. To test whether BON2-mediated lipid engagement underlies DolP’s distribution, we first examined the localization of DolP variants with reduced cardiolipin-binding ability. Consistent with the previous study by Bryant et al.^2^, DolP^W127E^ lost mid-cell localization in most cells, with residual enrichment only in some deeply constricted cells (Fig. 3A). Abolishing the cardiolipin-binding by removing the BON2 domain (DolP^ΔBON2^) diminished septal enrichment (Fig. 3B).

**Figure 3.**
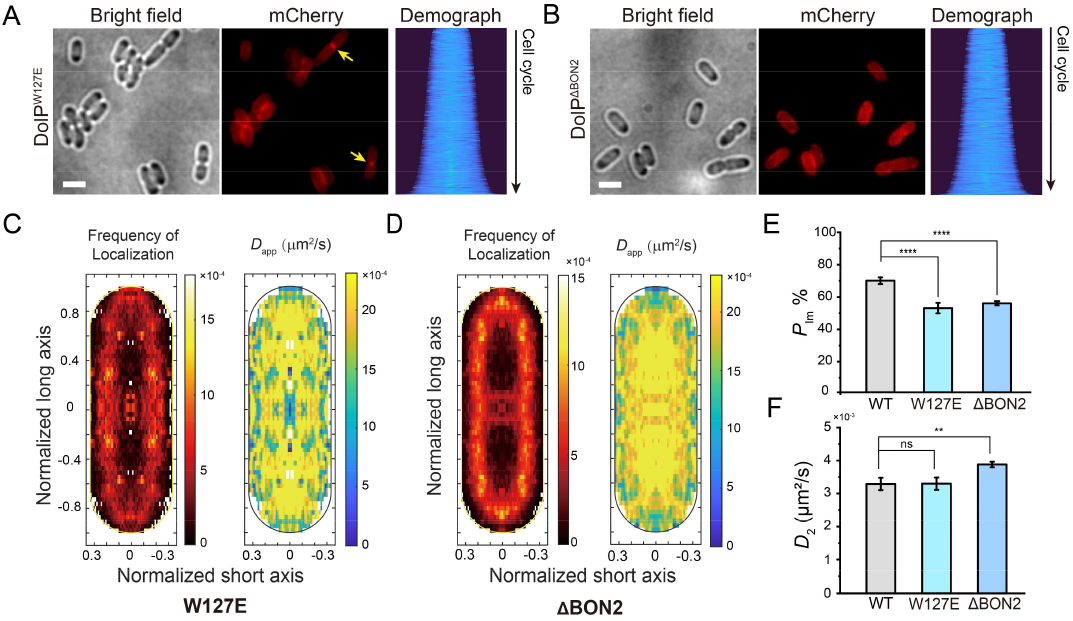
BON2-mediated cardiolipin binding governs DolP’s diffusion switch and mid-cell enrichment. **(A)** DolP^W127E^ (cardiolipin-binding site mutant) loses most mid-cell enrichment except for some deeply constricted cells (arrows). **(B)** DolP^ΔBON2^ (cardiolipin-binding domain deletion) abolishes mid-cell enrichment. Demographs at right in (A,B). Scale bars, 2 µm. More than 1000 cells from three independent experiments for each condition (Supplementary table 3). **(C**,**D)** Heatmaps of localization frequency (left) and *D*_app_ heatmaps (right) for W127E and ΔBON2 variants show loss of mid-cell pattern and increased mobility. **(E)** Immobile-state fraction is reduced in W127E and ΔBON2 variants. **(F)** Diffusive-state *D*_2_ is unchanged in W127E but increased in ΔBON2. ****P < 0.0001; **P < 0.01; ns, not significant (P > 0.05); two-tailed Student’s t-test. More than 270 trajectories from three independent experiments for each condition (Supplementary table 5).

To investigate whether DolP’s motion depends on the cardiolipin-binding domain, we performed spt-PALM to track PAmCherry fused DolP variant molecules. The localization heatmaps confirmed a lack of mid-cell accumulation for both variants (Fig. 3C,D). Mobility state analysis revealed how BON2 tunes DolP mobility. Relative to wildtype DolP, the immobile state of DolP^W127E^ significantly decreased (from 69 ±2% to 52 ±3%; Fig. 3E, Supplementary Table 5), suggesting reduced residency in the immobile state with reduced cardiolipin-binding ability. Interestingly, a similar *D*_2_ value was observed for the diffusive state (*D*_2_ = 0.0033 ±0.0002 µm^2.^s^−1^), indicating that the diffusive state is an intrinsic property of DolP, not a consequence of cardiolipin binding (Fig. 3F, Supplementary Table 5). The reduced fraction of immobile DolP^ΔBON2^ trajectories also supported the result. However, the diffusive state of DolP^ΔBON2^ showed a slightly higher diffusion coefficient (*D*_2_ = 0.0039 ±0.0001 µm^2.^s^−1^), indicating that BON2 deletion both weakens lipid/protein engagement and alters DolP’s hydrodynamic property (e.g., molecule size/conformation).

As an upper-bound control for minimal membrane engagement, replacing both BON1 and BON2 with PAmCherry yielded a much faster lipoprotein reporter (DolP^ΔBON12^) requiring 200-ms exposure to capture most moving molecules. This variant showed no mid-cell localization (Fig. S3A) and elevated average diffusion coefficient (<*D*_linear_>= 0.0076 ±0.0003 µm^2.^s^−1^; Fig. S3B), on the same order of magnitude of another OM lipoprotein PAL without its PG binding-domain^26^. CDF analysis still revealed two mobility states, one fast-diffusion state (*D*_2_ = 0.013 ±0.001 µm^2.^s^−1^, 51 ±2%) and another slow-diffusion state (*D*_1_ = 0.0014 ±0.0003 µm^2.^s^−1^, 49 ±2%; Fig. S3C, Supplementary Table 5), pointing to an intrinsic heterogenous inner-leaflet environment of the OM. The difference from DolP^ΔBON2^ also indicate that BON1 contributes to the slow diffusion of DolP in an unknown manner. Together, this mutant series demonstrates that BON2-dependent cardiolipin binding slows DolP and traps it at the constriction site, explaining its septal enrichment, which may occur without the need for a stable association with the divisome.

### Active constriction is required for DolP recruitment, independent of EnvC/AmiAB

If curvature-driven cardiolipin enrichment is the proximate cue for DolP trapping, this model predicts that divisome assembly alone would be insufficient, and that active septal synthesis (causing constriction) must be required. We therefore inhibited septal PG transpeptidation with aztreonam (FtsI-specific β-lactam), which does not disassemble the early/late divisome proteins immediately but halts septal formation and cell constriction. As expected, FtsN still accumulated at mid-cell after a 30-min aztreonam treatment (Fig. S4), but DolP redistributed largely to the lateral wall, with residual mid-cell localization only in cells that had already shown envelope invagination (Fig. 4A,B). Meanwhile, single DolP localizations lost mid-cell enrichment and the immobile fraction (Fig. 4C–E), paralleling BON2/W127E behaviors. Diffusive-state *D*_2_ increased modestly (*D*_2_ = 0.0041 ±0.0001 µm^2.^s^−1^; Supplementary Table 5), likely due to the decreased integrity of OM upon β-lactam treatment^27^. Because FtsW was also shown accumulated at the division site after aztreonam treatment^18^, our data indicate that ongoing constriction, rather than the mid-cell presence of FtsWI or FtsN, creates the DolP-trapping environment, likely via cardiolipin-enriched membranes.

**Figure 4.**
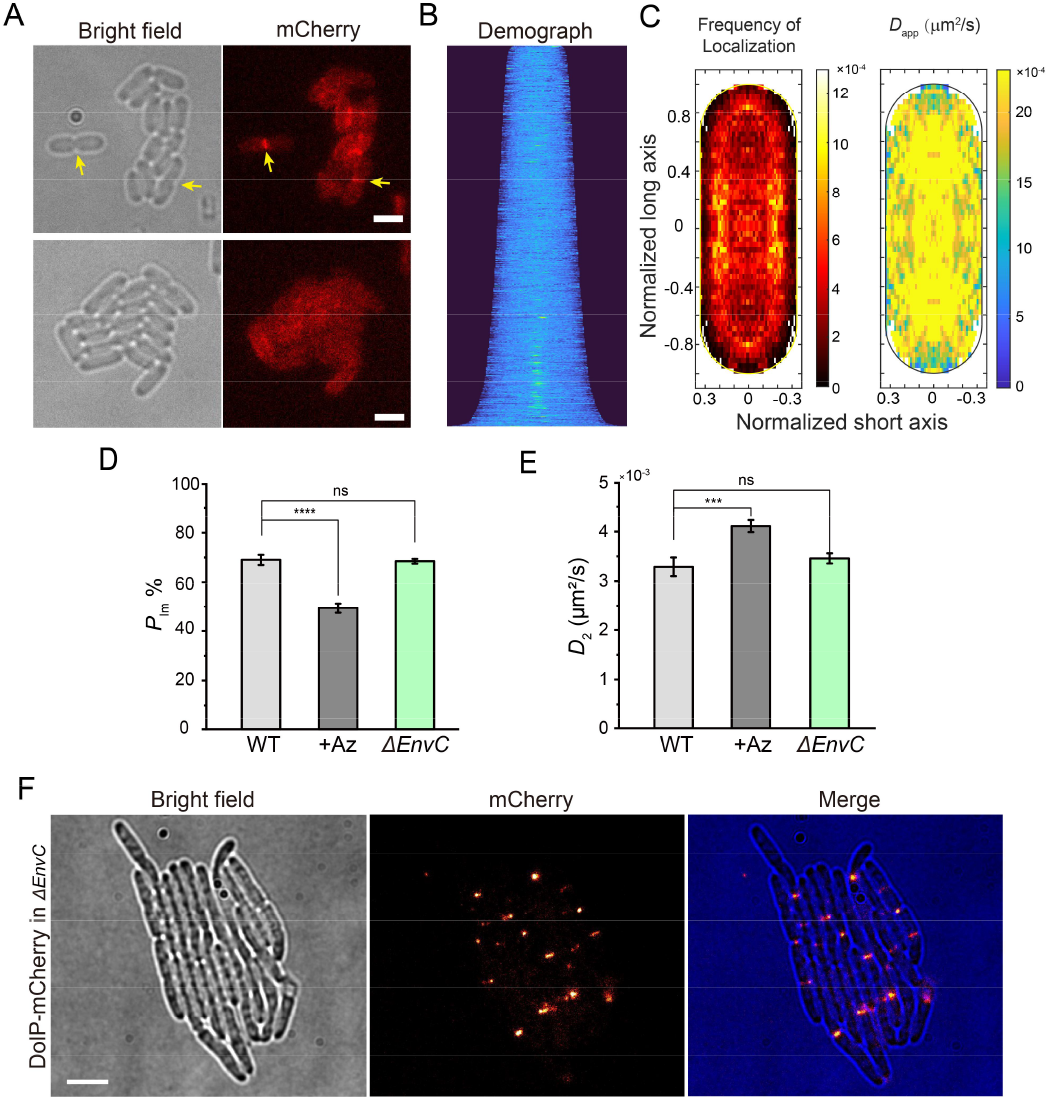
Active constriction is required for DolP trapping, and EnvC– AmiAB is dispensable. **(A)** Aztreonam (FtsI inhibitor) treatment halts septal constriction and redistributes DolP–mCherry to the lateral envelope (arrows mark residual septal signal in pre-constricted cells). Scale bars, 2 µm. **(B)** Demograph after aztreonam shows reduced mid-cell enrichment. More than 1000 cells from three independent experiments for each condition (Supplementary table 3). **(C)** Localization and *D*_app_ heatmaps confirm loss of mid-cell and slow-diffusion trajectories under aztreonam. **(D**,**E)** spt-PALM quantification: aztreonam reduces the immobile fraction (∼49%) and increases diffusive-state *D*_2_; *envC* deletion (Δ*envC*) leaves both metrics unchanged (ns). ****P < 0.0001; ***P < 0.001; ns, not significant (P > 0.05); two-tailed Student’s t-test. More than 430 trajectories from three independent experiments for each condition (Supplementary table 5). **(F)** In elongated Δ*envC* cells, DolP remains concentrated at constriction sites. Scale bars, 4 µm.

Moreover, we asked whether DolP enrichment is related to EnvC because DolP was implicated in cell separation parallel to the EnvC-AmiAB pathway. In Δ*envC* strain, DolP still accumulated at division sites (Fig. 4F), and spt-PALM showed no significant changes in immobile fraction or diffusive-state *D*_2_ (Fig. 4D,E), indicating that DolP’s localization and diffusion state switching are independent of EnvC. We explain that even in the absence of the EnvC–AmiAB pathway, the NlpD-activated AmiC (which can partially compensate for this loss)^5^ enables sufficient constriction to generate high-curvature, cardiolipin-enriched zones that recruit DolP.

## Discussion

How the lipoprotein DolP is recruited to the division site in diderm bacteria to orchestrate outer membrane constriction has remained elusive. Here, we show that DolP occupies two mobility states and is recruited to constricting septa via a cardiolipin-dependent shift toward the immobile state, rather than through stable association with late divisome proteins (FtsWI, FtsN) or dependence on EnvC. Thus, DolP likely senses the negative curvature of the constricting septum by binding cardiolipin, providing an OM-side mechanism that coordinates OM invagination with cytokinesis.

Spt-PALM analysis quantitively characterized two types of movement of DolP in the OM: an immobile state with *D*_1_ ∼ 1.0×10^−4^ µm^2.^s^−1^ and a diffusive state with *D*_2_ ∼ 3.3×10^−3^ µm^2.^s^−1^. The diffusion coefficient of the mobile DolP state is comparable to that of other outer membrane proteins (OMPs), such as OmpF^28^. As observed in some OMPs, a ∼0.5-μm region was found to confine DolP’s diffusion, indicating the existence of micro-domains on the inner leaflet of OM. The immobile state commonly found in OMPs or OM lipoproteins while they are covalently linked to PG. For instance, PAL, another cell division related OM lipoprotein, also exhibits extensive immobility unless its PG binding site is blocked^26^. Given no evidence showing DolP linked to PG but bound with cardiolipin, we reason that the local enrichment of cardiolipin trap and immobilize the movement of DolP. Indeed, DolP variants that mutated or truncated the cardiolipin-binding domain, BON2, diffuse faster with a decrease of the immobile population. The rest BON1 might still interact with other unknown factors, because removing this domain together with BON2 resulted in fast mobility, similar to other OM lipoproteins not bound to PG (PAL in *E. coli* and PapS in *Rhodospirillum rubrum*)^26,29^.

As cardiolipin is known to prefer the highly curved regions on the membrane, such as cell poles and the division site^25^, it naturally serves as a spatial cue for DolP’s (immobile state) accumulation. In this work, we found the immobile state of DolP molecules enrich at the mid-cell while the W127E and ΔBON2 variants lost this mid-cell pattern, demonstrating that the cardiolipin-mediated diffusion-state switch to immobile state is the key to recruit DolP to the division site. Moreover, our results also suggest that DolP is unlikely trapped in the division site via direct interaction with the septal PG synthesis or hydrolysis proteins.

It remains unclear what other patterners affect the mobility of DolP (e.g., with BON1). One recent study showed that the core OMP synthesis protein BamA can interact with DolP, which might contribute to the residual immobile state in our experiments. While DolP’s function in division is still unresolved^5,15^, a critical future goal is to define its relationship with the NlpD-AmiC pathway.

## Methods

### Bacterial strains and culture conditions

*E. coli* DH5α cells were used for plasmid subcloning, while *E. coli* BW25113 cells served as the parental strain for phenotypic assays and imaging experiments. *E. coli* strains were cultured in Luria–Bertani (LB) medium (1% tryptone, 0.5% yeast extract, 1% NaCl) at the indicated temperatures, with 1.5% agar added for solid media. Unless otherwise specified, both liquid and solid cultures were maintained at 37°C. When necessary, antibiotics were supplemented at the following concentrations: kanamycin, 50 μg/mL; carbenicillin, 50 μg/mL; chloramphenicol, 75 μg/mL. For fluorescence imaging and spt-PALM, cells were grown and incubated in M9 medium (1× M9 salts, 0.4% glucose, 1× MEM amino acids, 2 mM MgSO_4_, 0.1 mM CaCl_2_) at 25°C. For protein induction, arabinose (Ara) (Sigma, #E003256) or anhydrotetracycline (aTC) (Shanghai yuanye, #S25906) was added at the concentrations indicated, induction concentration for each experiment is listed in Supplementary Table 3,4.

### Strain construction

#### ΔdolP/envC strains

Strains were constructed by λ-Red recombination in *E. coli* BW25113. A kanamycin-resistance (Kan^R^) cassette was amplified from pKD13^30^ with Phanta Max Super-Fidelity DNA Polymerase (Vazyme, #P505-d1) using primers oJC121 and oJC122, each carrying a 50-nt extension homologous to sequences flanking *dolP*^31^.The PCR product was electroporated into BW25113 cells bearing pKD46^30^, and recombinants were selected on LB agar containing 50 μg/mL kanamycin and validated by Sanger sequencing, yielding strain JC123 (BW25113, *dolP::aph*). The kanamycin marker was subsequently excised by FLP recombinase expressed from pCP20^32^, generating strain JC125 (BW25113, *dol<>frt*). Deletion of *envC* was performed similarly, yielding strain JC144 (BW25113, *envC::aph*).

#### ΔenvCΔdolP strain

The Kan^R^ cassette was transferred into strain JC125 bearing pKD46, and recombinants were selected on LB agar containing 50 μg/mL kanamycin. Positive clones were validated by colony PCR and sequencing, yielding strain JC145 (BW25113, *dolP<>frt, envC::aph*).

### Plasmids construction

Plasmid pJC32 was generated to express DolP-mNeonGreen (linker: GGSSLVPSSDP)^33^ under the control of an aTC-inducible pLtetO-1 promoter^34^. The plasmid backbone containing the mNeonGreen encoding gene was amplified from pYY071 using primers oJC91 and oJC95. The *dolP* gene fragment was amplified from *E. coli* strain BW25113 genomic DNA with primers oJC96 and oJC97. The fragments were assembled by seamless cloning using the ClonExpress method (Vazyme, ClonExpress II One Step Cloning Kit, #C116).

Plasmid pJC42 was generated to express wildtype DolP. The plasmid backbone was amplified from pJC32 using primers oJC136 and oJC137 and ligated to remove the fragment of mNeonGreen encoding gene using the ClonExpress method.

Plasmid pJC23 was generated as an empty vector control. The plasmid backbone was amplified from pZH509 (a gift from Dr. Hensel^34^) using primers oJC90 and oJC94, and the product was circularized using the ClonExpress method.

Plasmids pJC39 and pJC40, were generated to express DolP fused to mCherry or PAmCherry, respectively, via a polypeptide linker (GGSSLVPSSDP) under the control of an aTC-inducible promoter. Plasmids were constructed using the same procedure as described for pJC32. The plasmid backbone was amplified using primers oJC102 and oJC103. The mCherry fragment was amplified from pJWK1^35^ using primer pair oJC101 and oJC125. The PAmCherry fragment was amplified from pYH002^36^ using primer pair oJC126 and oJC127. The fragments were ligated by seamless cloning using the ClonExpress method.

Plasmids pJC49 and pJC50 carrying a single-residue substitution DolP^W127E^ fused to mCherry or PAmCherry, respectively, were generated by site-directed mutagenesis using primers oJC143 and oJC144, and the product was circularized using the ClonExpress method.

Plasmids encoding fusion proteins DolP^ΔBON2^-mCherry (pJC54), DolP^ΔBON2^-PAmCherry (pJC57), DolP^ΔBON12^-mCherry (pJC55) and DolP^ΔBON12^-PAmCherry (pJC58) were constructed based on pJC39 and pJC40 using the same seamless cloning method. The primers used for plasmid construction are listed in Supplementary Table 2.

### Sample preparation for light microscopy imaging

#### Bacterial cell preparation

*E. coli* cells were recovered from glycerol stocks by streaking onto LB agar plates containing the appropriate antibiotics and incubating at 37°C for 24 h. Single colonies were subsequently inoculated into M9 medium and cultured overnight at 25°C and allowed to reach log-phase (OD_600_ = 0.2–0.6). For the aztreonam-treated condition, cells were incubated with 2 μg/mL aztreonam for 30 min prior to harvest.

#### Agarose gel pad preparation

For imaging, 2% agarose gel pads (LONZA, 50111) were prepared by dissolving 100 mg of low-melting agarose in 500 μL of M9 medium. The mixture was heated at 70°C for 1 h until fully dissolved and subsequently maintained at 50°C until use. Antibiotics were supplemented when required^37^. Approximately 80 μL of the molten agarose was applied onto a clean glass slide and sandwiched between a clean coverslip and a 0.5 mm thick rubber gasket (Bioptechs). The pads were allowed to solidify at room temperature (25°C) for a minimum of 2 h. ∼0.8 μL cell culture were deposited onto the solidified pad and subsequently covered with a clean coverslip. For the aztreonam-treated condition, aztreonam was pre-mixed in the agarose gel to reach a final concentration of 2 μg/mL.

### Light microscopy

HiLO (highly inclined and laminated optical sheet) imaging^38^ was conducted on a Nikon ECLIPSE Ti2 inverted microscope equipped with a 100× oil-immersion TIRF objective (NA = 1.49) and either a BSI or Prime95B sCMOS camera (Teledyne Photometrics) as previously described^39^. The HiLO illumination mode was implemented and adjusted using an MTW150 TIRF illuminator (Beijing Coolight Technology). Excitation was supplied by a solid-state laser engine (MULT4, Beijing Coolight Technology) producing continuous-wave lasers at 405 nm, 488 nm and 561 nm, which were applied for mCherry and mNeonGreen excitation as well as for the photoactivation and excitation of PAmCherry. Fluorescence emission was collected via a ZT405/488/561/647 rpc dichroic mirror (Chroma Technology).

For two-color and single-color imaging, the optical system was operated with an additional 1.5 × magnification and used the BSI camera as detector, yielding an effective pixel size of 43.3 nm. Two-color images were acquired sequentially with 488 nm and 561 nm excitation, and brightfield images were recorded as references. The specific imaging conditions are shown in Supplementary Table 3.

For single-particle tracking PALM (spt-PALM), the Prime95B camera was used (sensitivity mode), resulting in an effective pixel size of 110 nm. Fluorescence emission in the 561 nm channel was collected through an ET605/50m emission filter (Chroma Technology). Single DolP-PAmCherry molecules were photoactivated using a 405 nm laser, and were subsequently tracked under continuous 561 nm laser illumination. The imaging sequence was automatically acquired by the Cellvision Software (Beijing Coolight Technology). The specific imaging conditions are shown in Supplementary Table 4.

### Light microscopy data processing

#### Cell length measurements and localization analysis

Bright-field images of cells were segmented using Cellpose3.0^40^, and both segmentation results and raw images were imported into MicrobeJ^41^ plugin in Fiji^42^ to extract the outlines and medial axes. The morphological features were measured, including cell length and fluorescence intensity profiles along the long axis, with more than 80 cells analyzed per strain. For localization analysis and demograph construction, medial-axis fluorescence profiles were imported to MATLAB, where profiles were normalized, cells were ordered by length, and demographs were generated accordingly. The timing of protein localization at the mid-cell was assessed by extracting, smoothing, and fitting fluorescence profiles at the cell center, and recording the normalized cell-cycle position corresponding to 50% of the maximum intensity.

#### spt-PALM

2D-single-molecule tracking were performed as previously described^18^. Briefly, localizations were identified using ThunderSTORM^43^ in Fiji and subsequently linked into trajectories using a maximum distance threshold of 300 nm per frame and a dark interval threshold of 5 frames. Additionally, only trajectories last longer than 5 frames were retained for analysis. Extracellular trajectories were excluded based on bright-field images and Cellpose3.0 segmented results. Trajectory datasets from multiple fields of view were then combined for the calculation of MSD and CDF.

#### MSD

The mean squared displacement (MSD) of individual trajectories was calculated using a custom MATLAB script as previously described^44^. Trajectories were first filtered to a minimum length of 5 or 11 frames (Supplementary Table 4) and the mean squared displacements at each *Δt* were calculated with the error from 1000 times of bootstrapping.

The MSD curves were further analyzed using linear or Kusumi fitting approaches. For linear fitting, the first four points of each MSD curve were fitted to:

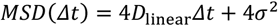

where *D*_linear_ annotates the diffusion coefficient assuming the molecules with free Brownian motion. *σ* is related to the blurry from localization error and the motion within a single exposure. This parameter is also used in the CDF fitting later.

For confined diffusion, the first ten points of the MSD curve were fitted using the Kusumi model^45^:

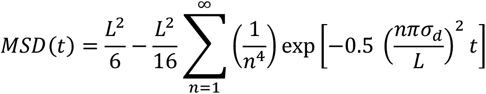

where *L* is the confinement length and *σ*_d_ is the displacement parameter. Experimental MSD curves were fitted using nonlinear least-squares optimization to extract *L* and *σ*_d_. The confinement length was also related to the MSD plateau value according to:

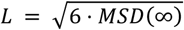

The corrected diffusion coefficient was calculated as:

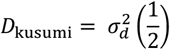

To estimate the error of the *D*_linear_ and *L*, the MSD results were bootstrapped 1000 times and fit with the corresponding equation.

#### CDF

Trajectories with a minimum length of 5 or 11 frames (Supplementary Table 4) were retained without an upper length limit. For each trajectory, the squared displacement *r*^2^ was calculated by measuring the vector displacement between consecutive frames at a lag time of Δ*t*=exposure time. Cumulative radial distribution function (CDF) analysis of single step squared displacements were fitted using a unified single or two-population diffusion model as previously described^46^.

For a single diffusion population, the CDF is given by:

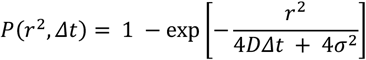

For two distinct diffusion populations, the CDF can be expressed as:

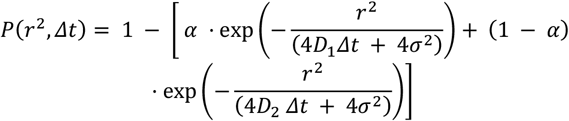

In the displacement analysis, *r*^2^ is the squared displacement between consecutive frames at lag time Δ*t*, and *P* (*r*^2^, Δ*t*) is the cumulative probability of displacements ≤ *r*^2^ across all trajectories. *D*_1_ and *D*_2_ are the diffusion coefficients of the two mobility states, and *α* represents the fraction of particles in state 1. σ^2^ was calculated from the MSD fitting above.

To estimate the error of the diffusion coefficients and population percentage, the CDF curves were bootstrapped 1000 times and fit with the corresponding equation (double-population).

#### Localization and diffusion heatmap

To quantify spatial organization of localization and diffusion state, the coordinates of individual localizations were recalculated refereeing to a normalized cell (3:1, length: width), using custom MATLAB scripts. Briefly, cell contours were first segmented by Cellpose3.0 using the bright field images. The main axis (long axis) of each contour was extracted by a PCA analysis and aligned horizontally. For each localization of a trajectory, the distances to the long and short axis were calculated and normalized by the cell length and width, generating a pair of coordinates [x, y] in a normalized cell frame. Single-step diffusion coefficients (*D*_raw_) were calculated for each pair of consecutive localizations. The coefficients were calibrated using the relation 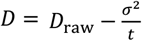 accounting for measurement error. Each calibrated diffusion coefficient was assigned to the mean position between two consecutive localizations. To exploit the elliptical symmetry and enhance spatial sampling, trajectory coordinates were first converted to absolute values, confining all points to the first quadrant (x ≥ 0, y ≥ 0). Each first-quadrant point was then symmetrically projected to the remaining three quadrants, effectively quadrupling the sampling density while preserving spatial relationships. The cell area was discretized into a 55×55 grid, and for each bin, the average diffusion coefficient 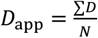 and data point percentage 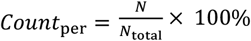 were computed and represented as heatmaps.

## End Matter

### Author Contributions

J.X. and X.Y. conceptualized the study. J.X. performed the experiments. J.X., D.Y., J.W., and X.Y. analyzed the data. Y. H. provided critical strains and methods. J.X. and X.Y. wrote the original draft. J.X., D.Y., J.W., Y. H., and X.Y. reviewed and edited the manuscript. X.Y. supervised the project. Funding was obtained by Y. H. and X.Y.

## Acknowledgments

We thank members of Yang’s laboratory for valuable discussions, and Jie Hu for guidance in molecular cloning and microscopy techniques. We also appreciate Dr. Zach Hensel, Dr. Hui Wang, Yan Zhang, Haofeng He, Yu Yan, and Yuhan Nie for helpful suggestions and providing plasmids used in this study. This work was supported by the National Natural Science Foundation of China (32270035 to X.Y., 32170032 and 32370034 to Y.H.), Project of MOE Key Laboratory of Geriatric Diseases and Immunology (No. KJS2601 to X.Y.), the Fundamental Research Funds for the Central Universities (WK9100000063 to X.Y.), and USTC start-up funding (KY9100000035, KJ2070000083 to X.Y.), Qilu Medical Foundation of Suzhou Medical College (SYQL2012 to Y.H.), and Jiangsu Province Health Innovation Team (Y.H.).

## Declaration of interests

The authors declare no competing financial interest.

## Resource availability

### Lead contact

Information and requests for reagents may be directed to, and will be fulfilled by, the lead contact, Xinxing Yang.

### Materials availability

Strains, plasmids, and probes are available on request from the lead contact with a material transfer agreement.

### Declaration of interests

The authors declare no competing financial interest.

## Notes

### Competing Interest Statement

The authors have declared no competing interest.

### Summary of Updates

I have edit the order of the author.

## Reference

1 Rojas, E. R. et al. The outer membrane is an essential load-bearing element in Gram-negative bacteria. Nature 559, 617–621 (2018). 10.1038/s41586-018-0344-3

2 Bryant, J. A. et al. Structure of dual BON-domain protein DolP identifies phospholipid binding as a new mechanism for protein localisation. Elife 9 (2020). 10.7554/eLife.62614

3 Deghelt, M. et al. Peptidoglycan–outer membrane attachment generates periplasmic pressure to prevent lysis in Gram-negative bacteria. Nature Microbiology 10, 1963–1974 (2025). 10.1038/s41564-025-02058-9

4 Egan, A. J. F. Bacterial outer membrane constriction. Mol Microbiol 107, 676–687 (2018). 10.1111/mmi.13908

5 Tsang, M. J., Yakhnina, A. A. & Bernhardt, T. G. NlpD links cell wall remodeling and outer membrane invagination during cytokinesis in Escherichia coli. PLoS Genet 13, e1006888 (2017). 10.1371/journal.pgen.1006888

6 Coltharp, C., Buss, J., Plumer, T. M. & Xiao, J. Defining the rate-limiting processes of bacterial cytokinesis. Proc Natl Acad Sci U S A 113, E1044–1053 (2016). 10.1073/pnas.1514296113

7 Cowles, C. E., Li, Y., Semmelhack, M. F., Cristea, I. M. & Silhavy, T. J. The free and bound forms of Lpp occupy distinct subcellular locations in Escherichia coli. Mol Microbiol 79, 1168–1181 (2011). 10.1111/j.1365-2958.2011.07539.x

8 Cascales, E., Bernadac, A., Gavioli, M., Lazzaroni, J. C. & Lloubes, R. Pal lipoprotein of Escherichia coli plays a major role in outer membrane integrity. J Bacteriol 184, 754–759 (2002). 10.1128/jb.184.3.754-759.2002

9 Lloubès, R. et al. The Tol-Pal proteins of the Escherichia coli cell envelope: an energized system required for outer membrane integrity? Research in Microbiology 152, 523–529 (2001). 10.1016/s0923-2508(01)01226-8

10 Heidrich, C. et al. Involvement of N-acetylmuramyl-L-alanine amidases in cell separation and antibiotic-induced autolysis of Escherichia coli. Mol Microbiol 41, 167–178 (2001). 10.1046/j.1365-2958.2001.02499.x

11 Bernhardt, T. G. & de Boer, P. A. The Escherichia coli amidase AmiC is a periplasmic septal ring component exported via the twin-arginine transport pathway. Mol Microbiol 48, 1171–1182 (2003). 10.1046/j.1365-2958.2003.03511.x

12 Peters, N. T., Dinh, T. & Bernhardt, T. G. A fail-safe mechanism in the septal ring assembly pathway generated by the sequential recruitment of cell separation amidases and their activators. J Bacteriol 193, 4973–4983 (2011). 10.1128/jb.00316-11

13 Navarro, P. P. et al. Cell wall synthesis and remodelling dynamics determine division site architecture and cell shape in Escherichia coli. Nat Microbiol 7, 1621–1634 (2022). 10.1038/s41564-022-01210-z

14 Lyu, Z. et al. Third track model for coordination of septal peptidoglycan synthesis and degradation by FtsN in Escherichia coli. Nat Microbiol 10, 1521–1534 (2025). 10.1038/s41564-025-02011-w

15 Boelter, G. et al. The lipoprotein DolP affects cell separation in Escherichia coli, but not as an upstream regulator of NlpD. Microbiology 168 (2022). 10.1099/mic.0.001197

16 Sun, S. & Chen, J. Unveiling the role of BON domain-containing proteins in antibiotic resistance. Front Microbiol 15, 1518045 (2024). 10.3389/fmicb.2024.1518045

17 Shaner, N. C. et al. Improved monomeric red, orange and yellow fluorescent proteins derived from Discosoma sp. red fluorescent protein. Nat Biotechnol 22, 1567–1572 (2004). 10.1038/nbt1037

18 Yang, X. et al. A two-track model for the spatiotemporal coordination of bacterial septal cell wall synthesis revealed by single-molecule imaging of FtsW. Nat Microbiol 6, 584–593 (2021). 10.1038/s41564-020-00853-0

19 Lyu, Z. et al. FtsN maintains active septal cell wall synthesis by forming a processive complex with the septum-specific peptidoglycan synthases in E. coli. Nat Commun 13, 5751 (2022). 10.1038/s41467-022-33404-8

20 Du, S. & Lutkenhaus, J. Assembly and activation of the Escherichia coli divisome. Mol Microbiol 105, 177–187 (2017). 10.1111/mmi.13696

21 Britton, B. M. et al. Conformational changes in the essential E. coli septal cell wall synthesis complex suggest an activation mechanism. Nat Commun 14, 4585 (2023). 10.1038/s41467-023-39921-4

22 Subach, F. V. et al. Photoactivatable mCherry for high-resolution two-color fluorescence microscopy. Nat Methods 6, 153–159 (2009). 10.1038/nmeth.1298

23 Yang, X. et al. GTPase activity-coupled treadmilling of the bacterial tubulin FtsZ organizes septal cell wall synthesis. Science 355, 744–747 (2017). 10.1126/science.aak9995

24 Bisson-Filho, A. W. et al. Treadmilling by FtsZ filaments drives peptidoglycan synthesis and bacterial cell division. Science 355, 739–743 (2017). 10.1126/science.aak9973

25 Renner, L. D. & Weibel, D. B. Cardiolipin microdomains localize to negatively curved regions of Escherichia coli membranes. Proc Natl Acad Sci U S A 108, 6264–6269 (2011). 10.1073/pnas.1015757108

26 Szczepaniak, J. et al. The lipoprotein Pal stabilises the bacterial outer membrane during constriction by a mobilisation-and-capture mechanism. Nature Communications 11, 1305 (2020). 10.1038/s41467-020-15083-5

27 Torabian, P. et al. The effect of clinically relevant betalactam, aminoglycoside, and quinolone antibiotics on bacterial extracellular vesicle release from E. coli. bioRxiv (2023). 10.1101/2023.11.22.568081

28 Spector, J. et al. Mobility of BtuB and OmpF in the Escherichia coli outer membrane: implications for dynamic formation of a translocon complex. Biophys J 99, 3880–3886 (2010). 10.1016/j.bpj.2010.10.029

29 Pöhl, S. et al. An outer membrane porin-lipoprotein complex modulates elongasome movement to establish cell curvature in Rhodospirillum rubrum. Nature Communications 15, 7616 (2024). 10.1038/s41467-024-51790-z

30 Datsenko, K. A. & Wanner, B. L. One-step inactivation of chromosomal genes in Escherichia coli K-12 using PCR products. Proc Natl Acad Sci 97, 6640–6645 (2000). 10.1073/pnas.120163297

31 Baba, T. et al. Construction of Escherichia coli K-12 inframe, single-gene knockout mutants: the Keio collection. Molecular Systems Biology 2 (2006). 10.1038/msb4100050

32 Cherepanov, P. P. & Wackernagel, W. Gene disruption in Escherichia coli: TcR and KmR cassettes with the option of Flp-catalyzed excision of the antibiotic-resistance determinant. Gene 158, 9–14 (1995). 10.1016/0378-1119(95)00193-a

33 Ranava, D. et al. Lipoprotein DolP supports proper folding of BamA in the bacterial outer membrane promoting fitness upon envelope stress. Elife 10, e67817 (2021). 10.7554/eLife.67817

34 Hensel, Z. A plasmid-based Escherichia coli gene expression system with cell-to-cell variation below the extrinsic noise limit. PLoS One 12, e0187259 (2017). 10.1371/journal.pone.0187259

35 Wang, J. et al. Class A PBPs reinforce the septal cell wall following initial synthesis by SEDS-bPBP pairs during bacterial cytokinesis. bioRxiv (2025). 10.1101/2025.09.12.675754

36 Nie, Y., Hu, J., Zhang, S., Meng, X. & Fu, G. Molecular resolution imaging based on two-color single-molecular localization microscopy (SMLM). Opt Express 33, 20023–20036 (2025). 10.1364/OE.559116

37 Ding, L. H. et al. Fluorogenic Probes for Real-Time Tracking of Bacterial Cell Wall Dynamics with Nanoscopy. Acs Nano 19, 14389–14403 (2025). 10.1021/acsnano.5c01930

38 Tokunaga, M., Imamoto, N. & Sakata-Sogawa, K. Highly inclined thin illumination enables clear single-molecule imaging in cells. Nat Methods 5, 159–161 (2008). 10.1038/nmeth1171

39 Yan, D., Xue, J., Xiao, J., Lyu, Z. & Yang, X. Protocol for single-molecule labeling and tracking of bacterial cell division proteins. STAR Protocols 5, 102766 (2024). 10.1016/j.xpro.2023.102766

40 Stringer, C. & Pachitariu, M. Cellpose3: one-click image restoration for improved cellular segmentation. Nat Methods 22 (2025). 10.1038/s41592-025-02595-5

41 Ducret, A., Quardokus, E. M. & Brun, Y. V. MicrobeJ, a tool for high throughput bacterial cell detection and quantitative analysis. Nat Microbiol 1, 16077 (2016). 10.1038/nmicrobiol.2016.77

42 Schindelin, J. et al. Fiji: an open-source platform for biological-image analysis. Nat Methods 9, 676–682 (2012). 10.1038/Nmeth.2019

43 Ovesny, M., Krizek, P., Borkovec, J., Svindrych, Z. K. & Hagen, G. M. ThunderSTORM: a comprehensive ImageJ plug-in for PALM and STORM data analysis and super-resolution imaging. Bioinformatics 30, 2389–2390 (2014). 10.1093/bioinformatics/btu202

44 Bettridge, K., Harris, F. E., Yehya, N. & Xiao, J. RNAP Promoter Search and Transcription Kinetics in Live E. coli Cells. The Journal of Physical Chemistry B 127, 3816–3828 (2023). 10.1021/acs.jpcb.2c09142

45 Kusumi, A., Sako, Y. & Yamamoto, M. Confined Lateral Diffusion of Membrane-Receptors as Studied by Single-Particle Tracking (Nanovid Microscopy) - Effects of Calcium-Induced Differentiation in Cultured Epithelial-Cells. Biophys J 65, 2021–2040 (1993). 10.1016/S0006-3495(93)81253-0

46 Törk, L., Moffatt, C. B., Bernhardt, T. G., Garner, E. C. & Kahne, D. Single-molecule dynamics show a transient lipopolysaccharide transport bridge. Nature 623, 814–819 (2023). 10.1038/s41586-023-06709-x

