## Supplementary Material for "Recruitment of the outer-membrane lipoprotein DolP to the division site via cardiolipin-mediated diffusion-state switching"

**This file includes:**

**Figures S1 to S4**

**Supplementary Table 1 to 6**

**References**

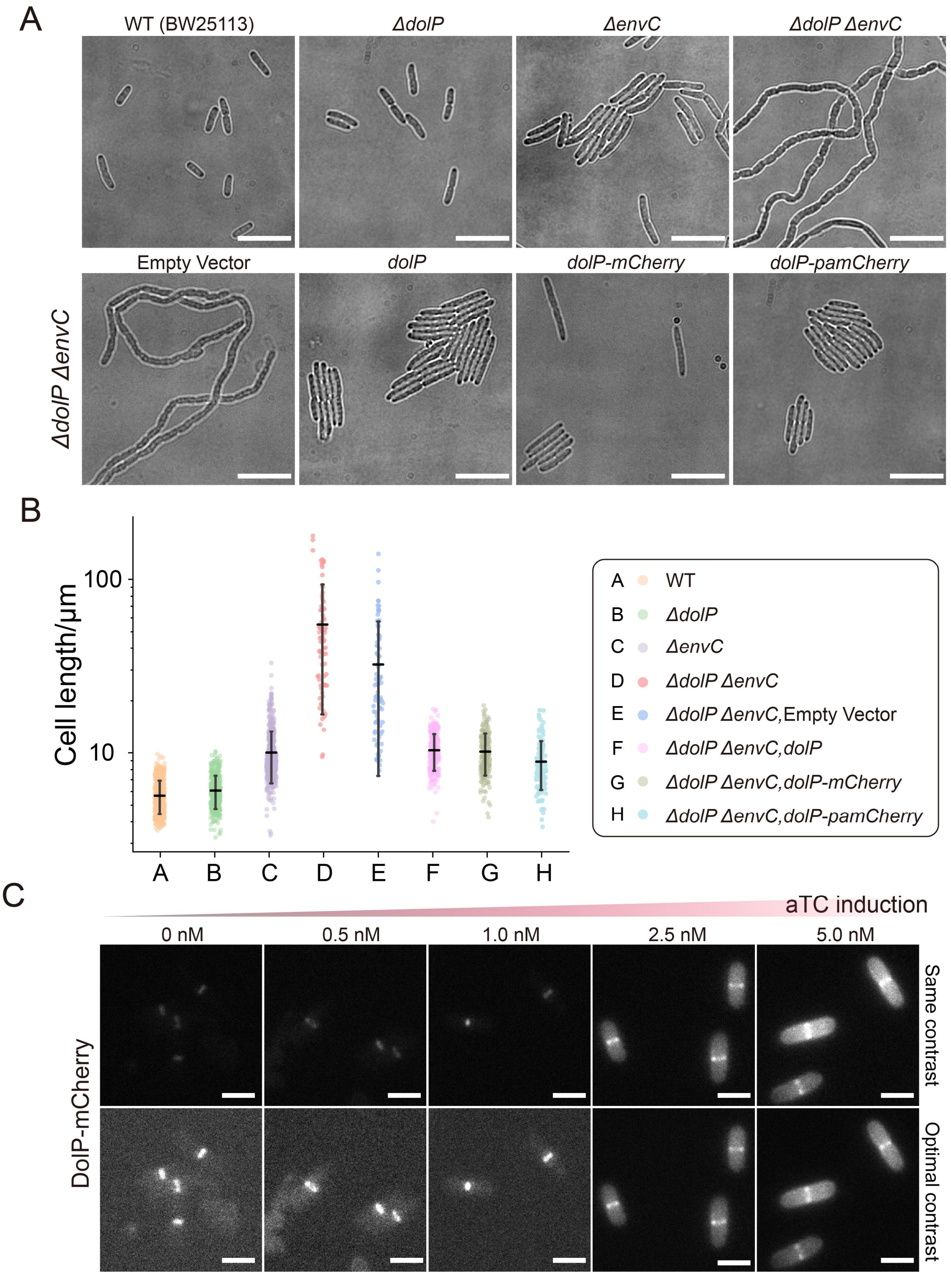
A
WT (BW25113)
ΔdolP
ΔenvC
ΔdolP ΔenvC
Empty Vector
dolP
dolP-mCherry
dolP-pamCherry
ΔdolP ΔenvC
B
Cell length/μm
100
10
A B C D E F G H
A WT
B ΔdolP
C ΔenvC
D ΔdolP ΔenvC
E ΔdolP ΔenvC,Empty Vector
F ΔdolP ΔenvC,dolP
G ΔdolP ΔenvC,dolP-mCherry
H ΔdolP ΔenvC,dolP-pamCherry
C
aTC induction
0 nM
0.5 nM
1.0 nM
2.5 nM
5.0 nM
DolP-mCherry
Same contrast
Optimal contrast

**Figure S1**. **DolP deletion causes chain formation in Δ*envC* mutants while DolP and its FP fusions rescues the phenotype; DolP localizes to the mid-cell independent of induction.**

**(A)** Representative bright field cell images of WT(BW25113), Δ*dolp,* Δ*envC,* Δ*dolP*Δ*envC* cells (up). Bright-field images of Δ*dolP*Δ*envC* strain harboring pJC23($P_{LtetO\text{-}1}$), pJC42($P_{LtetO\text{-}1}$::*dolP*), pJC39($P_{LtetO\text{-}1}$::*dolP-mCherry*), pJC40($P_{LtetO\text{-}1}$::*dolP-PAmCherry*) (bottom). All cells were grown in LB medium without aTC induction at 30˚C to $OD_{600}$ = 0.2–0.6. Scale bars, 10 µm.

**(B)** Cell length distribution of the strains shown in (A). Number of cells analyzed: WT, 645; Δ*dolP*, 418; Δ*envC*, 679; Δ*dolP*Δ*en*vC, 80; Δ*dolP*Δ*envC* empty vector, 86; Δ*dolP*Δ*envC*, *dolP*, 278; Δ*dolP*Δ*envC*, *dolP-mCherry*, 196; Δ*dolP*Δ*envC*, *dolP-pamCherry*, 137. Data are presented as mean ± s.d. from three independent replications.

**(C)** Representative images of Δ*dolP* strain harboring pJC39 ($P_{LtetO\text{-}1}$::*dolP-mCherry*). The contrast of each image was adjusted identically to allow comparison of fluorescence intensity at different aTC concentration (up). The contrast of each image was individually optimized to highlight the mid-cell localization of the protein (bottom). Cells were grown in M9-glucose medium with 0 nM, 0.5 nM, 1.0 nM, 2.5 nM or 5.0 nM aTC at 25˚C to $OD_{600}$ = 0.2–0.6. Scale bars, 2 µm.

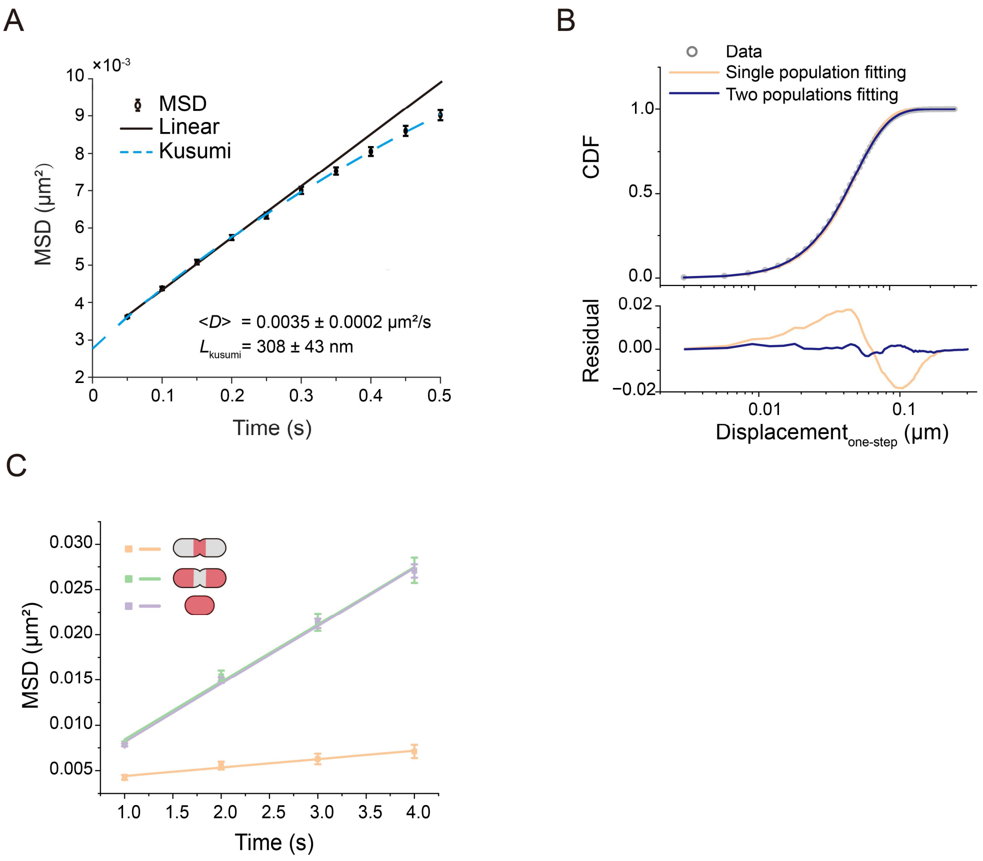

**Figure S2. Fast spt-PALM also revealed two diffusion populations of DolP.**

**(A)** MSD of DolP-PAmCherry molecules in Δ*dolp* cells with 50-ms exposure time (black circles) was fit by the linear function of the first four points (solid line) and Kusumi mode of the first ten points (dashed). Error bars indicate s.e.m. from bootstrapping, yielding $\langle D_{linear} \rangle$ = 0.0035 ±0.0002 μm²·s⁻¹ with confinement length $L_{kusumi}$ = 308 ±43 nm (mean ± s.e.m.). The standard error of the mean of $D_{linear}$ and $L_{kusumi}$ were obtained from 1000 bootstrapping resamples.

**(B)** CDF curve of DolP-PAmCherry molecules in Δ*dolp* cells with 50-ms exposure time (gray circles) was fitted by a double (blue curve) and a single (tan curve) population, with residuals indicated below.

**(C)** MSDs of DolP-PAmCherry molecules in different groups (as in Fig. 2H) (squares) were fitted by the linear function of the first four points (group I, purple line; group II, green line; group III, tan line). Error bars indicate s.e.m. from bootstrapping.

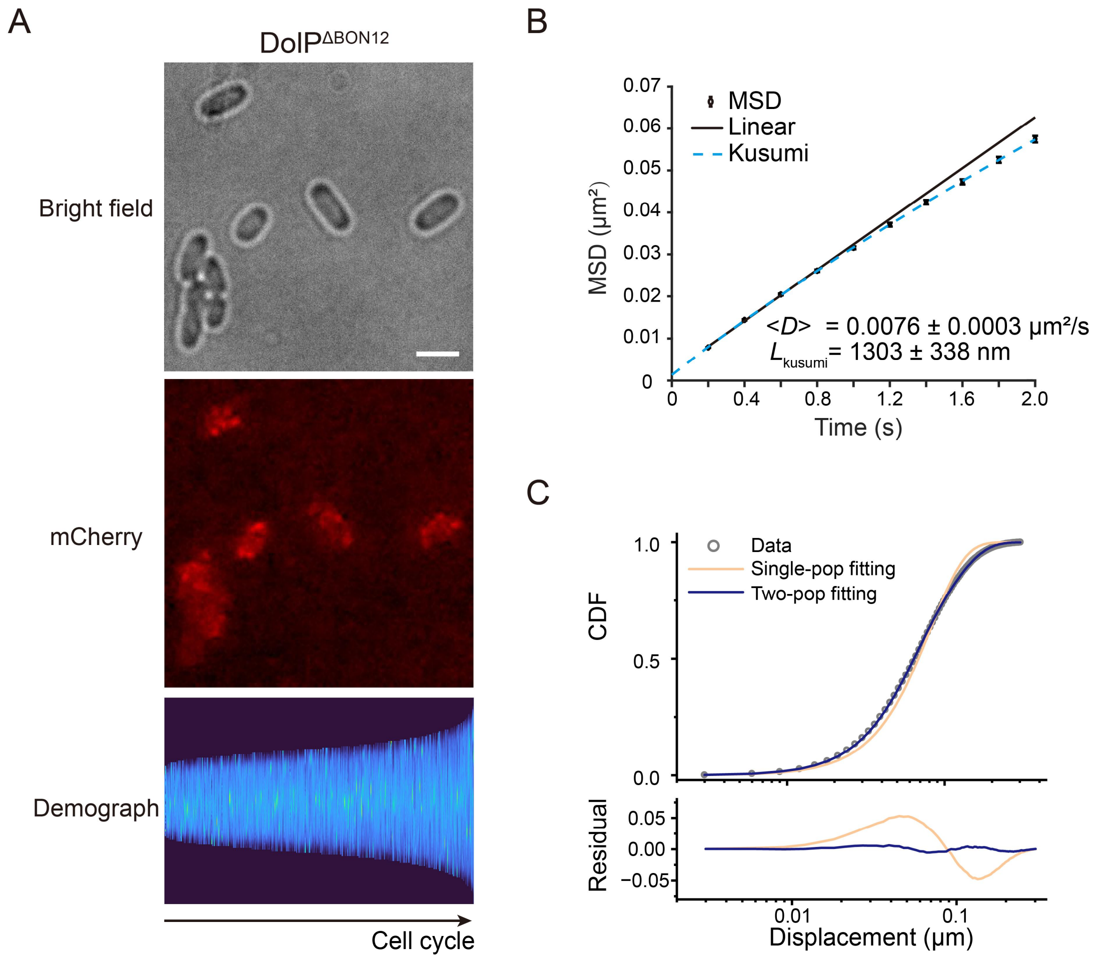

**Figure S3. Removal of both BON domains disrupts DolP's mid-cell localization and accelerates its diffusion.**

**(A)** Representative images of Δ*dolP* strain harboring pJC55 ($P_{LtetO\text{-}1}$:: *dolP*$^{\Delta BON12}$-*mCherry*) shows the loss of mid-cell enrichment, demonstrated in the fluorescence image(middle) and demograph (bottom). Scale bar, 2 μm. More than 1000 cells from three independent experiment for each condition (Supplementary table 3).

**(B)** MSD curve of DolP$^{\Delta BON12}$-PAmCherry in Δ*dolP* cells with 200-ms exposure time (black circles) was fit by the linear function of the first four points (solid line) and Kusumi model of the first ten points (dashed). Error bars indicate s.e.m. from bootstrapping, yielding $\langle D_{linear} \rangle = 0.0076 \pm 0.0003$ μm²·s⁻¹ with confinement length $L_{kusumi} = 1303 \pm 338$ nm (mean ± s.e.m.). The standard error of the mean of $D_{linear}$ and $L_{kusumi}$ were obtained from 1000 bootstrapping resamples.

**(C)** CDF curve of DolP$^{\Delta BON12}$ -PAmCherry molecules in Δ*dolP* cells with 200-ms exposure time (gray circles) was best fitted by a double (blue curve) instead of a single (tan curve) population, as indicated by the residuals below.

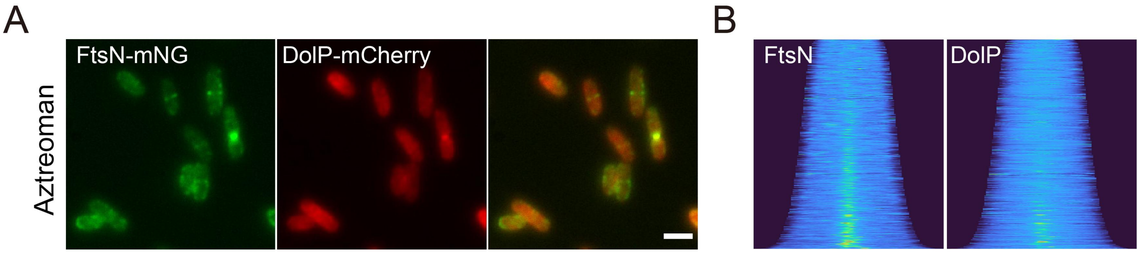

**Figure S4. DolP mid-cell localization requires active septal synthesis.**

**(A)** Two-color imaging of cells expressing DolP–mCherry and FtsN–mNeonGreen (late divisome) after aztreonam treatment. FtsN localizes at the mid-cell, whereas DolP does not. Scale bars, 2 µm. More than 1000 cells from three independent experiment for each condition (Supplementary table 3).

**(B)** Demographs of FtsN (left) and DolP (right) after aztreonam treatment, showing reduced mid-cell enrichment of DolP but FtsN does not, comparing to Fig 1B.

**Supplementary Table 1: Strains and plasmids used in the study.**

| Bacteria | Genotype | Reference/source |
|---|---|---|
| DH5α | *Δ(argF-lac)169, φ80dlacZ58(M15), ΔphoA8, recA1, endA1, thiE1, hsdR17, λ-, deoR481, gyrA96(NalR)* | CGSC #1423[1] |
| BW25113 | *Δ(araD-araB)567, ΔlacZ4787(::rrnB-3), rph-1, Δ(rhaD-rhaB)568, hsdR514* | CGSC #8590[2] |
| JC123 | BW25113 *dolP*::*aph* | This study |
| JC125 | BW25113 *dolP*::*frt* | This study |
| JC136 | BW25113 *dolP*::*frt,* $P_{LtetO-1}$::*dolP-mCherry* | This study |
| JC137 | BW25113 *dolP*::*frt,* $P_{LtetO-1}$::*dolP-PAmCherry* | This study |
| JC139 | BW25113 *dolP*::*frt,* $P_{LtetO-1}$::*dolP-mCherry,* $P_{T5lac}$::*mNeonGreen-zapA* | This study |
| JC144 | BW25113 *envC*::*aph* | This study |
| JC145 | BW25113 *dolP*::*frt, envC*::*aph* | This study |
| JC147 | BW25113 *dolP*::*frt,* $P_{LtetO-1}$::*dolP-mCherry,* $P_{T5lac}$:::*ftsN(E60-E61)-mNeonGreen*$^{SW}$ | This study |
| JC153 | BW25113 *dolP*::*frt, envC*::*aph,* $P_{LtetO-1}$::*dolP* | This study |
| JC154 | BW25113 *dolP*::*frt, envC*::*aph,* $P_{LtetO-1}$::*dolP-PAmCherry* | This study |
| JC155 | BW25113 *dolP*::*frt, envC*::*aph,* $P_{LtetO-1}$:: | This study |
| JC156 | BW25113 *dolP*::*frt, envC*::*aph,* $P_{LtetO-1}$::*dolP-mCherry* | This study |
| JC170 | BW25113 *dolP*::*frt,* $P_{LtetO-1}$::*dolP*$^{W127E}$*-mCherry* | This study |
| JC171 | BW25113 *dolP*::*frt,* $P_{LtetO-1}$::*dolP*$^{W127E}$*-PAmCherry* | This study |
| JC189 | BW25113 *dolP*::*frt,* $P_{LtetO-1}$::*dolP*$^{\Delta BON2}$*-mCherry* | This study |
| JC190 | BW25113 *dolP*::*frt,* $P_{LtetO-1}$::*dolP*$^{\Delta BON12}$*-mCherry* | This study |
| JC192 | BW25113 *dolP*::*frt,* $P_{LtetO-1}$::*dolP*$^{\Delta BON2}$*-PAmCherry* | This study |
| JC193 | BW25113 *dolP*::*frt,* $P_{LtetO-1}$::*dolP* $^{\Delta BON12}$*-PAmCherry* | This study |
| **Plasmid** | **Genotype** | **Reference/source** |
| pKD13 | R6K, *bla frt::aph::frt* | [2] |
| pKD46 | pSC101, *bla repA*$^{ts}$, *araC* $P_{BAD}$::*gam bet exo* | [2] |
| pCP20 | pSC101, *bla cat repA*$^{ts}$, *λ.p R-FLP cI857* | [3] |
| pJC23 | p15A, *bla,* $P_{LtetO-1}$ | This study |
| pJC32 | p15A, *bla,* $P_{LtetO-1}$::*dolP-mNeonGreen* | This study |
| pJC39 | p15A, *bla,* $P_{LtetO-1}$::*dolP-mCherry* | This study |
| pJC40 | p15A, *bla,* $P_{LtetO-1}$::*dolP-PAmCherry* | This study |
| pJC42 | p15A, *bla,* $P_{LtetO-1}$::*dolP* | This study |
| pJC49 | p15A, *bla,* $P_{LtetO-1}$:: *dolP*$^{W127E}$*-mCherry* | This study |
| pJC50 | p15A, *bla,* $P_{LtetO-1}$:: *dolP*$^{W127E}$*-PAmCherry* | This study |
| pJC54 | p15A, *bla,* $P_{LtetO-1}$:: *dolP*$^{\Delta BON2}$*-mCherry* | This study |
| pJC55 | p15A, *bla,* $P_{LtetO-1}$:: *dolP*$^{\Delta BON12}$*-mCherry* | This study |
| pJC57 | p15A, *bla,* $P_{LtetO-1}$:: *dolP*$^{\Delta BON2}$*-PAmCherry* | This study |
| pJC58 | p15A, *bla,* $P_{LtetO-1}$:: *dolP*$^{\Delta BON12}$*-PAmCherry* | This study |
| pYH002 | p15A, *bla,* $P_{LtetO-1}$:: *ffDronpa-ftsZ-PAmCherry* | [4] |
| pXY677 | ColE1, *cat, lacI*$^{Q}$ $P_{T5lac}$::*mNeonGreen-zapA* | [5] |
| pJL108 | ColE1, *cat, lacI*$^{Q}$ $P_{T5lac}$:::*ftsN(E60-E61)- mNeonGreen*$^{SW}$ | [6] |

| | | |
|---|---|---|
| pJWK1 | pJWK, $P_{1261}$::*mCherry* | [7] |
| pYY071 | p15A, *bla,* $P_{LtetO-1}$:: *ftsZ-mNeonGreen* | Lab stock |
| pZH509 | p15A, *bla,* $P_{LtetO-1}$:: *gfpmut2* | [8] |

**Supplementary Table 2: DNA oligos used in the study.**

| Primer | Sequence (5' - 3') | Purpose |
|---|---|---|
| 1 | TGATCGATAACACGCTTTTCCCTCACCAGGATGATTAAGGAGAATACATGATTCCGGGGATCCGTCGACC | Amplification of *aph* for JC123 |
| 2 | AGCGTCGCATCAGGCATTACAAGGGGCTGCTATTTAATAAACGTAAACGCTGTAGGCTGGAGCTGCTTCG | |
| 3 | AAGAGATGACTGGTAAGCCGCTGTTCATCGTGGAATAATCCCTCCCCATGATTCCGGGGATCCGTCGACC | Amplification of *aph* for JC144 and JC145 |
| 4 | AGAACGTTACGACGAAATGGAAACAAAACTTATCTTCCCAACCACGGCTGTGTAGGCTGGAGCTGCTTCG | |
| 5 | GCGGTACCTTTCTCCTCTTT | Amplification of vector backbone for pJC32 |
| 6 | CTGGTGCCGAGCAGCGATCCGGTGAGCAAAGGCGAAGAAGAT | |
| 7 | AGAGGAGAAAGGTACCGCATGAAGGCATTATCGCCAATCGC | Amplification of the gene encoding DolP for pJC32 |
| 8 | CGGATCGCTGCTCGGCACCAGGCTGCTGCCGCCTTTAATAAACGTAAACGCCGTAGTTAC | |
| 9 | TTTAATAAACGTAAACGCCGTAGTTAC | Amplification of vector backbone for pJC42 |
| 10 | TTTACGTTTATTAAATAATCTAGCAGGAGGATTTCACC | |
| 11 | TCTAGCAGGAGGATTTCACC | Amplification of vector backbone for pJC23 |
| 12 | TGAAATCCTCCTGCTAGAGCGGTACCTTTCTCCTCTTT | |
| 13 | CGGATCGCTGCTCGGCAC | Amplification of vector backbone for pJC39 and pJC40 |
| 14 | TAATCTAGCAGGAGGATTTCA | |
| 15 | GGTGAAATCCTCCTGCTAGATTACTTGTACAGCTCGTCCATGC | Amplification of the gene encoding mCherry for pJC39 |
| 16 | TGGTGCCGAGCAGCGATCCGGTGAGCAAGGGCGAGGAG | |
| 17 | TGGTGCCGAGCAGCGATCCGGTAAGCAAAGGCGAAGAGGAC | Amplification of the gene encoding PAmCherry for pJC40 |
| 18 | GGTGAAATCCTCCTGCTAGATTACTTGTACAGTTCATCCATGCC | |
| 19 | GCATCTAACGATACGGAAATCACCACCAAAGTGCGTTCGC | Mutagenesis of *dolP* for pJC49 and pJC50 |
| 20 | CGTATCGTTAGATGCTTCGCCCA | |
| 21 | AGATGCTTCGCCCAGACCA | Mutagenesis of *dolP* for pJC54 and |

| | | |
|---|---|---|
| 22 | CTGGGCGAAGCATCTGGCGGCAGCAGCCTGG | pJC57 |
| 23 | CACCTGGGTGCCGACACT | Mutagenesis of *dolP* for pJC55 and pJC58 |
| 24 | GTCGGCACCCAGGTGGGCGGCAGCAGCCTGG | |

**Supplementary Table 3. Cell culture, imaging conditions, and statistics of each figure. (two-color and single-color imaging)**

| Figure | Strain | Culture | Imaging | Statistics |
|---|---|---|---|---|
| 1B,C-up | JC139 | No inducer | 488 nm, 193 W/cm$^2$, 50 ms;<br>561 nm, 195 W/cm$^2$, 50 ms.<br>**(HiLO, Excitation, Power density, exposure time)** | Three replicates;<br>Demograph:<br>1787 cells. |
| 1B,C-bottom | JC147 | No inducer | 488 nm, 193 W/cm$^2$, 50 ms;<br>561 nm, 195 W/cm$^2$, 50 ms.<br>**(HiLO, Excitation, Power density, exposure time)** | Three replicates;<br>Demograph:<br>1241 cells. |
| 3A | JC170 | 2.5 nM aTC | 561 nm, 195 W/cm$^2$, 50 ms.<br>**(HiLO, Excitation, Power density, exposure time)** | Three replicates;<br>Demograph:<br>1665 cells. |
| 3B | JC189 | 2.5 nM aTC | 561 nm, 195 W/cm$^2$, 50 ms.<br>**(HiLO, Excitation, Power density, exposure time)** | Three replicates;<br>Demograph:<br>1313 cells. |
| 4A | JC136 | 1.0 nM aTC;<br>+Aztreonam | 561 nm, 39 W/cm$^2$, 50 ms.<br>**(HiLO, Excitation, Power density, exposure time)** | Three replicates;<br>Demograph:<br>1338 cells. |
| 4F | JC156 | No inducer | 561 nm, 39 W/cm$^2$, 50 ms.<br>**(HiLO, Excitation, Power density, exposure time)** | One replicate. |
| S1C | JC136 | 0/1.0/2.5/5.0 nM aTC | 561 nm, 39 W/cm$^2$, 50 ms.<br>**(HiLO, Excitation, Power density, exposure time)** | One replicate. |
| S3A | JC190 | No inducer | 561 nm, 195 W/cm$^2$, 50 ms.<br>**(HiLO, Excitation, Power density, exposure time)** | Three replicates;<br>Demograph:<br>1539 cells. |
| S4A | JC147 | No inducer;<br>+Aztreonam | 488 nm, 193 W/cm$^2$, 50 ms;<br>561 nm, 195 W/cm$^2$, 50 ms.<br>**(HiLO, Excitation, Power density, exposure time)** | Three replicates;<br>Demograph:<br>1167 cells. |

**Supplementary Table 4. Cell culture, imaging conditions, and statistics of each figure. (spt-PALM)**

| Protein | Figure | Strain | Culture | Imaging | Trajectory length |
|---|---|---|---|---|---|
| $DolP^{WT}$ | 1E,2A-E | JC137 | 2.5 nM aTC | 405 nm, 0.05 W/cm$^2$, 5 s;<br>561 nm, 6 W/cm$^2$, 1 s<br>**(HiLO, Excitation, Power density, exposure time)** | ≥11 frames |
| $DolP^{WT}$ | 2F-J, S2C | JC137 | 2.5 nM aTC | 405 nm, 0.05 W/cm$^2$, 5 s;<br>561 nm, 6 W/cm$^2$, 1 s<br>**(HiLO, Excitation, Power density, exposure time)** | ≥5 frames |
| $DolP^{W127E}$ | 3C,D | JC171 | 2.5 nM aTC | 405 nm, 0.05 W/cm$^2$, 5 s;<br>561 nm, 6 W/cm$^2$, 1 s<br>**(HiLO, Excitation, Power density, exposure time)** | ≥ 5 frames |
| $DolP^{W127E}$ | 3E,F | JC171 | 2.5 nM aTC | 405 nm, 0.05 W/cm$^2$, 5 s;<br>561 nm, 6 W/cm$^2$, 1 s<br>**(HiLO, Excitation, Power density, exposure time)** | ≥11 frames |
| $DolP^{\Delta BON2}$ | 3C,D | JC192 | 2.5 nM aTC | 405 nm, 0.05 W/cm$^2$, 5 s;<br>561 nm, 6 W/cm$^2$, 1 s<br>**(HiLO, Excitation, Power density, exposure time)** | ≥5 frames |
| $DolP^{\Delta BON2}$ | 3E,F | JC192 | 2.5 nM aTC | 405 nm, 0.05 W/cm$^2$, 5 s;<br>561 nm, 6 W/cm$^2$, 1 s<br>**(HiLO, Excitation, Power density, exposure time)** | ≥11 frames |
| $DolP^{WT}$ | 4C | JC137 | 2.5 nM aTC;<br>+Aztreonam | 405 nm, 0.05 W/cm$^2$, 5 s;<br>561 nm, 6 W/cm$^2$, 1 s<br>**(HiLO, Excitation, Power density, exposure time)** | ≥5 frames |
| $DolP^{WT}$ | 4D,E | JC137 | 2.5 nM aTC;<br>+Aztreonam | 405 nm, 0.05 W/cm$^2$, 5 s;<br>561 nm, 6 W/cm$^2$, 1 s<br>**(HiLO, Excitation, Power density, exposure time)** | ≥11 frames |
| $DolP^{WT}$<br>(Δ*envC)* | 4D,E | JC154 | 2.5 nM aTC | 405 nm, 0.05 W/cm$^2$, 5 s;<br>561 nm, 6 W/cm$^2$, 1 s<br>**(HiLO, Excitation, Power density, exposure time)** | ≥11 frames |
| $DolP^{WT}$ | S2A,B | JC137 | 2.5 nM aTC | 405 nm, 0.05 W/cm$^2$, 5 s;<br>561 nm, 39 W/cm$^2$, 50 ms<br>**(HiLO, Excitation, Power density, exposure time)** | ≥11 frames |

| | | | | | |
|---|---|---|---|---|---|
| DolP$^{\Delta BON12}$ | S3B,C | JC193 | No inducer | 405 nm, 0.5 W/cm$^2$, 5 s;<br>561 nm, 13 W/cm$^2$, 200 ms<br>**(HiLO, Excitation, Power density, exposure time)** | ≥11 frames |

**Supplementary Table 5. DolP dynamics under different conditions. (≥11 frames)**

| Group | $D_{linear}$ (μm²/s) | $D_{kusumi}$ (μm²/s) | $L_{kusumi}$ (nm) | $D_1$ (μm²/s) | $D_2$ (μm²/s) | Immobile Fraction $\alpha$ | Number of trajectories |
|---|---|---|---|---|---|---|---|
| WT | 0.0012 ±0.0001[a] | 0.0033 ±0.0004 | 676 ±346 | 0.000087 ±0.000028 | 0.0033 ±0.0002 | 0.69 ±0.02 | 432 |
| W127E | 0.0017 ±0.0001 | 0.0038 ±0.0005 | 1740 ±6830 | 0.000002 ±0.000044 | 0.0033 ±0.0002 | 0.52 ±0.03 | 272 |
| ΔBON2 | 0.0020 ±0.0001 | 0.0059 ±0.0003 | 854 ±57 | 0.00026 ±0.00002 | 0.0039 ±0.0001 | 0.55 ±0.01 | 1412 |
| +AZ | 0.0023 ±0.0001 | 0.0064 ±0.0005 | 981 ±104 | 0.00030 ±0.00003 | 0.0041 ±0.0001 | 0.49 ±0.02 | 605 |
| WT(Δ*envC)* | 0.0011 ±0.0001 | 0.0034 ±0.0002 | 651 ±48 | -0.00010[b] ±0.00001 | 0.0035 ±0.0001 | 0.69 ±0.01 | 2038 |
| ΔBON12-200ms | 0.0076 ±0.0003 | 0.018 ±0.001 | 1303 ±338 | 0.0014 ±0.0003 | 0.0127 ±0.0005 | 0.49[c] ±0.02 | 1398 |

[a.] The errors (s.e.m.) were estimated by calculated from 1000 bootstrapping.

[b.] The diffusion coefficient calibrated by the localization uncertainty could generate negative values, indicating nearly zero *D* or immobile behavior.

[c.] The percentage of the slow-diffusion state.

**Supplementary Table 6. DolP$^{WT}$ dynamics at different subcellular positions. (1-second exposure, ≥5 frames)**

| Group | $D_{linear}$ (μm²/s) | $D_{kusumi}$ (μm²/s) | $L_{kusumi}$ (nm) | $D_1$(μm²/s) | $D_2$ (μm²/s) | Immobile Fraction $\alpha$ | Number of trajectories |
|---|---|---|---|---|---|---|---|
| Group I | 0.0016 ±0.0001[a] | 0.0055 ±0.0006 | 632 ±77 | 0.000014 ±0.000037 | 0.0033 ±0.0001 | 0.58 ±0.02 | 843 |
| Group II | 0.0016 ±0.0003 | 0.0068 ±0.0016 | 572 ±113 | 0.000026 ±0.000035 | 0.0038 ±0.0003 | 0.62 ±0.04 | 260 |
| Group III | 0.00023 ±0.00008 | 0.00056 ±0.00109 | 246 ±7080 | -0.0000070[b] ±0.0000343 | 0.0030 ±0.0006 | 0.83 ±0.04 | 98 |

[a.] The errors (s.e.m.) were estimated by calculated from 1000 bootstrapping.

[b.] The diffusion coefficient calibrated by the localization uncertainty could generate negative values, indicating nearly zero $D$ or the immobile state.

**References:**

1 Chen, J., Li, Y., Zhang, K. & Wang, H. Whole-Genome Sequence of Phage-Resistant Strain Escherichia coli DH5α. *Genome Announc* **6** (2018). https://doi.org/10.1128/genomeA.00097-18

2 Datsenko, K. A. & Wanner, B. L. One-step inactivation of chromosomal genes in Escherichia coli K-12 using PCR products. *Proc Natl Acad Sci U S A* **97**, 6640-6645 (2000). https://doi.org/10.1073/pnas.120163297

3 Cherepanov, P. P. & Wackernagel, W. Gene disruption in Escherichia coli: TcR and KmR cassettes with the option of Flp-catalyzed excision of the antibiotic-resistance determinant. *Gene* **158**, 9-14 (1995). https://doi.org/10.1016/0378-1119(95)00193-a

4 Nie, Y., Hu, J., Zhang, S., Meng, X. & Fu, G. Molecular resolution imaging based on two-color single-molecular localization microscopy (SMLM). *Opt Express* **33**, 20023-20036 (2025). https://doi.org/10.1364/OE.559116

5 Yang, X. *et al.* A two-track model for the spatiotemporal coordination of bacterial septal cell wall synthesis revealed by single-molecule imaging of FtsW. *Nat Microbiol* **6**, 584-593 (2021). https://doi.org/10.1038/s41564-020-00853-0

6 Lyu, Z. *et al.* FtsN maintains active septal cell wall synthesis by forming a processive complex with the septum-specific peptidoglycan synthases in E. coli. *Nat Commun* **13**, 5751 (2022). https://doi.org/10.1038/s41467-022-33404-8

7 Wang, J. *et al.* Class A PBPs reinforce the septal cell wall following initial synthesis by SEDS-bPBP pairs during bacterial cytokinesis. *bioRxiv* (2025). https://doi.org/10.1101/2025.09.12.675754

8 Hensel, Z. A plasmid-based Escherichia coli gene expression system with cell-to-cell variation below the extrinsic noise limit. *PLoS One* **12**, e0187259 (2017). https://doi.org/10.1371/journal.pone.0187259
